# Contrasting effects of accumulation and tolerance characteristics in *Arabidopsis thaliana* under Cr(III) and Cr(VI) stress

**DOI:** 10.1101/2021.12.26.474207

**Authors:** Yonghong Han, Guotao Ding, Peng Sun, Giuiying Li, Weihao Li

## Abstract

In this study, for the first time we investigated Cr(III) and Cr(VI) stress-induced physiological and biochemical responses in *Arabidopsis thaliana*. The capacity of A. thalian to accumulate Cr is closely related to the valence of chromium. Cr(VI) was more toxic than Cr(III) as indicated by chromium accumulation and growth inhibition. When the concentration of chromium is greater than 200μM, the root length and biomass of A. thaliana are reduced. But interestingly, Cr(III) at 200μM increased the root length and biomass of A. thaliana compared to the control. The transmission electron microscope shows that Cr(VI) can cause the chloroplasts damaged and the chlorophyll reduced more than Cr(III). The chloroplasts were filled the starch grains. An increase of lipid peroxidation in A. thaliana roots caused by Cr was measured, and this effect increases as the increasing Cr. It indicated that A. thaliana suffers from Cr-induced oxidative stress which resulted cell death in roots. To fight against oxidative stress, Ascorbate peroxidase and Glutathione reductase were activated by Cr in antioxidant defense. The inhibition of growth, the accumulation of chromium, the responses of antioxidant systems, and the ultra-morphological changes indicate that Cr(VI) was more toxic than Cr(III).

## Introduction

Heavy metal is a major source of pollutants worldwide, leading to harmfully affecting human health when contaminants enter into agricultural lands and food chain.[1, 2]

Chromium (Cr) is widely used in many industrial fields such as electrodeposition, leather tanning, wood preservation, and pigments [3-5]. Due to the wide application of Chromium, large amounts of chromium-containing waste (liquid, solid and gaseous waste) are discharged into the environment, which eventually leads to biological and ecological pollution[6, 7]. The positive trivalent and the positive hexavalent are the most stable, although chromium can exhibit multiple forms (from −2 to+6). [8]

In recent years, more and more researchers pay close attention to the toxicity to plant by Cr. In most case, however, the toxicity data were about total Cr or Cr(VI) concentration[9, 10]. Research shows that different valence metals toxicity to aquatic organisms are not same. Therefore, in the risk assessment, biological toxicities which generated by the speciation of metals are indispensable.

The mobility, the bioavailability and the toxicity of Cr(III) and Cr(VI) are different. Cr(III) is an essential microelement necessary for the glucose metabolism in mammals, but not for the plants. In contrast, Cr(VI) is known carcinogens and classified as Group A (USEPA 1999). Previous studies generally believe that Cr(VI) has more toxic effects than Cr(III), but some studies have found the opposite result [11, 12]. These contradictory conclusions have aroused the interest of our research team.

To our knowledge, there are not any reports in which physiological and biochemical responses in *A. thaliana* was investigated under the stresses of Cr(III) and Cr(VI) [13, 14]. For this purpose, our team prepared medium containing different concentrations of Cr(III) and Cr(VI) and applied it to *A. thaliana* culture. In this study, we evaluated the effects of different concentrations of Cr(III) and Cr(VI)on the physiological biochemical and the enrichment effects of *A. thaliana*. Furthermore, evidence for altering the ultrastructural level of *A. thaliana* after chromium stress is also provided.

## Materials and methods

### Plant preparation

The seeds are washed with 75% ethanol once and washed with distilled water for 5 times (1 minute each time). The washed seeds were vernalized 3 days in the dark at 4 °C in petri dishes containing MS agar. *A. thaliana* grown in long-day conditions (16-h-light/8-h-dark photoperiod)90μM m^-2^ s^-1^ PAR, 70% RH, and 22°C. After 7 days, the plants were transferred to MS solution with different concentrations (from 100 to 700μM) of Cr(III) supplied as Cr(NO)_3_, or Cr(VI) supplied as K_2_Cr_2_O_7_. The control group was grown for 14 days without adding Cr. The continuously MS solution was renewed once a week.

### Growth status measurements

30 seedlings for each treatment were harvested after 14 days. The root length of the sample under different treatment was measured separately. To measure the dry mass, the sample is weighed after constant weight. The mean values were calculated.

### Determination of Cr Contents

Roots and leaves were harvested after 14 days of chromium stress, rinsed with 5% EDTA for one time, and rinsed 5 times with sterile water (5 minutes each time). Leaves and roots were pre-dried to a constant weight at 60 degrees Centigrade, and were ashed at 550°C. The ashing product was dissolved with 2 mL 1:1 (v/v) HNO_3_, transmit into 25 ml volumetric flask and bring to volume by 0.1% NaOH solution. [15, 16] The concentration of Cr was accurately measured by an atomic absorption spectrophotometer (HITACHI Z5000). Each sample was repeated five replicates.

### Measurement of Photosynthetic Pigments

After 14 days of chromium stress, 0.2g fresh leaves were extracted with 80% acetone in the dark by tissue homogenization method until the color of leaf tissue completely disappeared. The concentrations of chlorophyll (a, b) and carotenoids were detected using the spectrophotometer (723N, Jingke, Shanghai, China).

### Estimation of antioxidant enzymes and lipid peroxidation

4mL of 4°C pre-cooled 50 mM phosphate buffer (pH 7.8) were added into 1.0 g seedling roots in an ice bath. The sample containing the phosphate buffer was homogenized. The homogenate was centrifuged at 12000 rpm for 10 minutes at 4 °C. Discard the precipitate and the supernatant for subsequent experiments. The Lipid peroxidation increasing was estimated by malondialdehyde (MDA) method according to reported paper of Zhou et al[17]. The activity of superoxide dismutase (EC 1.15.1.1, SOD) was measured by the nitroblue tetrazolium (NBT) method[18]. Catalase (EC 1.11.1.6, CAT) activity was detected by the method of Aebi[19]. Peroxidase (EC 1.11.1.7, POD) activity was analyzed according to the method of Zhou[20]. Ascorbate peroxidase (EC 1.11.1.11, APX) activity was measured by the method of Nakano and Asada[21]. Glutathione reductase (EC 1.6.4.2, GR) activity was analyzed by Garcia-Limones et al[22].

### Ultramicroscopic Observations and Trypan Blue staining

The method that we used to detect the dead tissues and cells is Trypan blue. According to the protocols of our senior scholars, Chen et al[23], 10 root samples under each treatment were treated for 15 minutes in 0.04% trypan blue aqueous solution. The stained samples were washed 4 times with sterile water (2 minutes each time), and were soaked overnight.

Under the chromium stress for 14 days, the leave samples were soaked with 2.5% glutaraldehyde for 12 hours at 4 °C. The fixed sample was rinsed 4 times with phosphate buffer (0.1 M, pH 7.0) for 5 minutes each time. Then, we used the 1% OsO4 phosphate buffer to fix samples for 2 hours, and after that the fixed samples were washed with phosphate buffer 4 times(5 minutes each time). Subsequently, the samples were dehydrated with different concentrations of ethanol, which the volume concentration of ethanol was 30%, 50%, 70%, 90%.. Finally the leaves were coated with Epon resin[24]. The Ultracut Eultramicrotome (Leica, Germany) was used for ultrathin sections of 70 nm thickness. The microstructure was observed by a JEL-1230 transmission electron microscope (TEM, Hitachi, Japan).

### Statistical analysis

Statistical analysis of the data was performed using the SPSS 16.0 (SPSS, Chicago, IL, USA). Each experiment was repeated at least three times. Different letters on tables and histograms indicate that a difference at the P < 0.05 level.

## Results and discussion

### Growth Inhibition of *A. thaliana* exposure to Cr(III) and Cr(VI)

**Fig 1. A** shows the morphology of three-week-old *A. thaliana* exposed to graded concentrations of Cr. It shows Cr concentrations in the culture medium of up to 200μM exhibited no obvious toxicity on seeding growth of *A. thaliana*, but exhibited significantly suppress gradually at the concentration of 300μM, 500μM and 700μM. As roots are the first one to come in contact with the Cr, their growth receive the most impact. In this study, the decrease of root length was observed from 300μM for Cr(III) and Cr(VI). The highest root length decrease was obtained for 700μM concentration with 51% and 57% for Cr(III) and Cr(VI), respectively(Fig 1 B). Interestingly, a relatively low concentration (200μM) of Cr(III) caused a significant increase in dry biomass as compared to the control. On the contrary, dry biomass decreased gradually from 300μM to 700μM for Cr(III) and Cr(VI). In the concentration of 700μM, the decrease was up to 47% and 73% for Cr(III) and Cr(VI), respectively(Fig 1 C).

**Fig 1.**
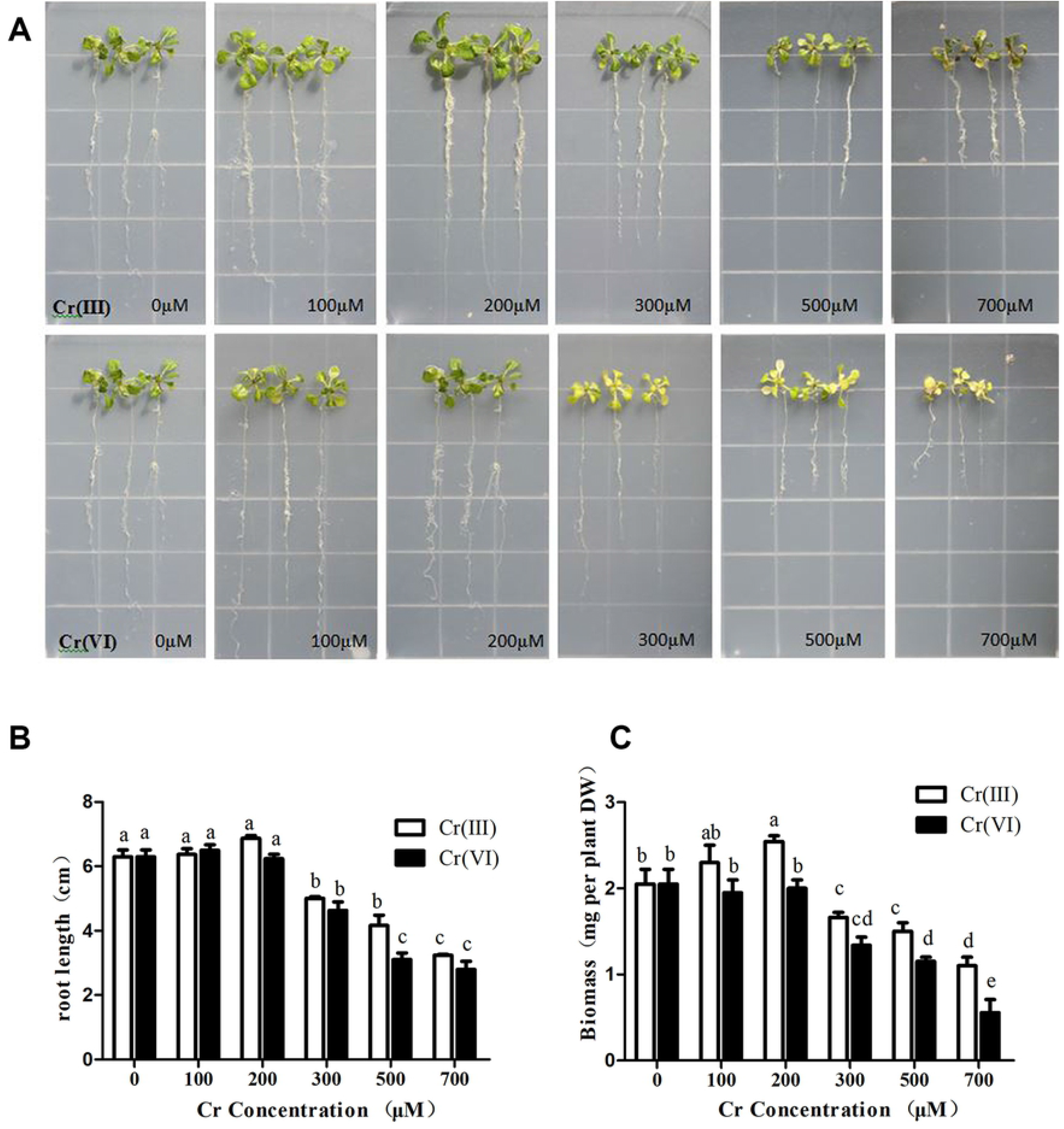
Physiological effects of Cr(III) and Cr(VI) on A. thaliana. (A) Images of A. thaliana exposed to Cr(III) and Cr(VI); (B) Root length; (C) Dry biomass. Data are mean ± standard error of five replicates. Different letters on histograms are statistically different (P < 0.05).

It is proved that the toxicity of Cr(VI) is stronger than that of Cr(III) by measuring the growth of A. thaliana. Whether Cr(III) or Cr(VI), when the concentration of Cr reaches or exceeds 200μM, the inhibition of A. thaliana growth was observed. But interestingly, at 200μM, Cr(III) caused a statistically significant increase in dry biomass as compared to the control. It indicated a relatively low concentration of Cr(III) can stimulate A. thaliana growth.

### Cr uptake and accumulation in *A. thaliana*

**Fig 2**. indicates the content of chromium in different parts of *A. thalian*. The capacity of *A. thalian* to accumulate Cr is closely related to the valence of chromium. The accumulation of Cr in roots, it do not distinguish between Cr(III) and Cr(VI), was higher than in leaves. After concentration were measured, the Cr(VI) was higher than the Cr(III) in root tissues. The concentration of chromium in roots can reach up to 1650μg g^-1^ DW under 700 mM Cr(VI) stress, whereas only 1129μg g^-1^ exposed to 700μM Cr(III).

**Fig. 2.**
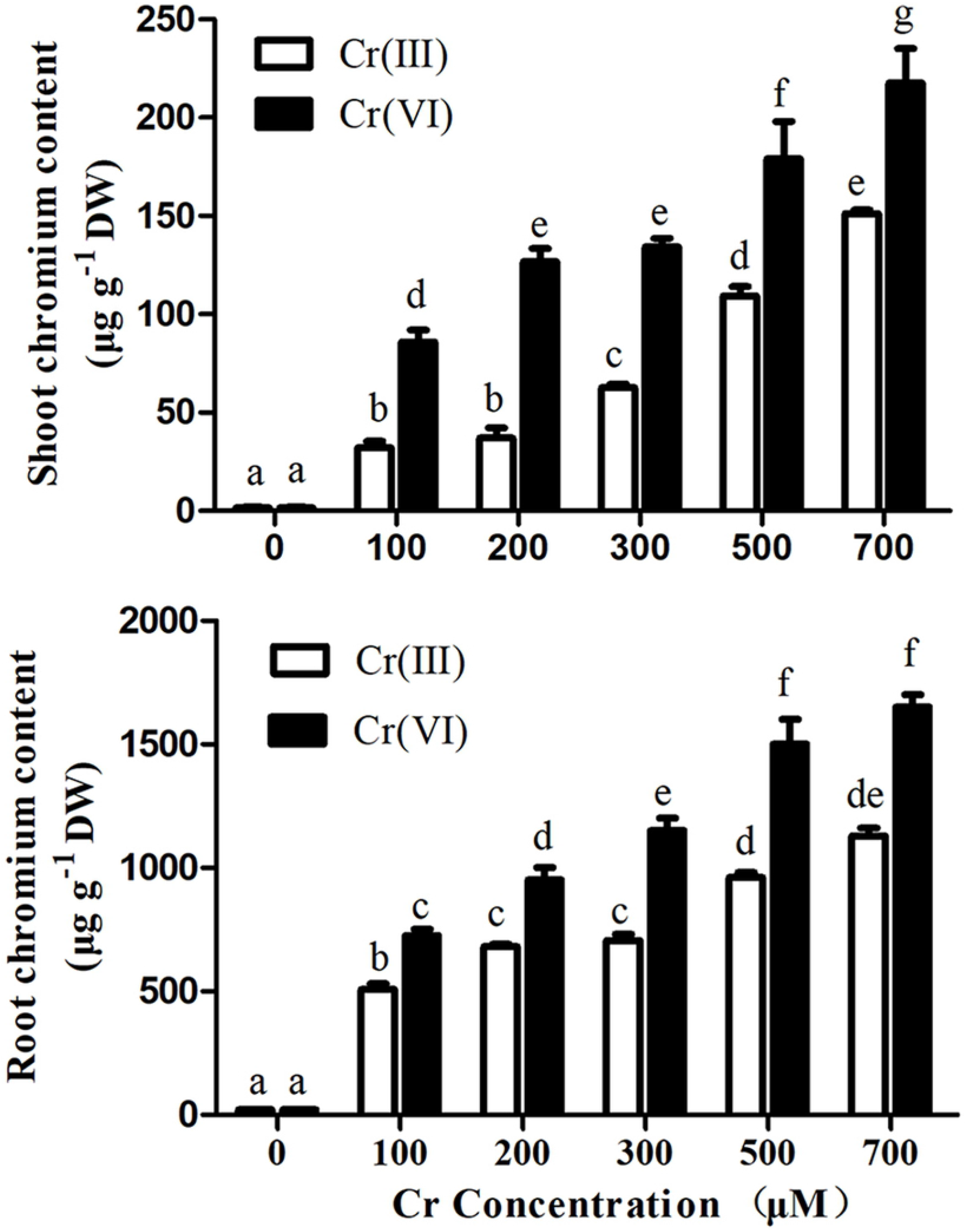
The contents of Cr in roots and shoots of A. thaliana, after treated by Cr(III) and Cr(VI)for 14 d. Data are mean ± standard error of five replicates. Different letters on histograms are statistically different (P < 0.05).

### Photosynthetic pigments analysis

Chlorophyll is a key index for the degree of toxicity by abiotic stresses on plants[25-27]. Under the chromium stress for 14 days, the total chlorophyll contents of fresh leaves were measured. The contents of chlorophyll a, chlorophyll b and total chlorophyll were reduced in *A. thalian*, under the stress of 100μM Cr(VI). Total chlorophyll fell by 53%, under the stress of Cr(VI) (Table 1). Moreover, the ratio of chlorophyll a/b obviously increase under the treatment of 200μM Cr(VI) indicated that chlorophyll b was more sensitive than the chlorophyll a under the stress. The maximum is 2.38. The effect of Cr(III)on chlorophyll of *A. thalian* is relatively light. The radio of chlorophyll a/b is reduced only with 700μM Cr(III).

**Table 1.**
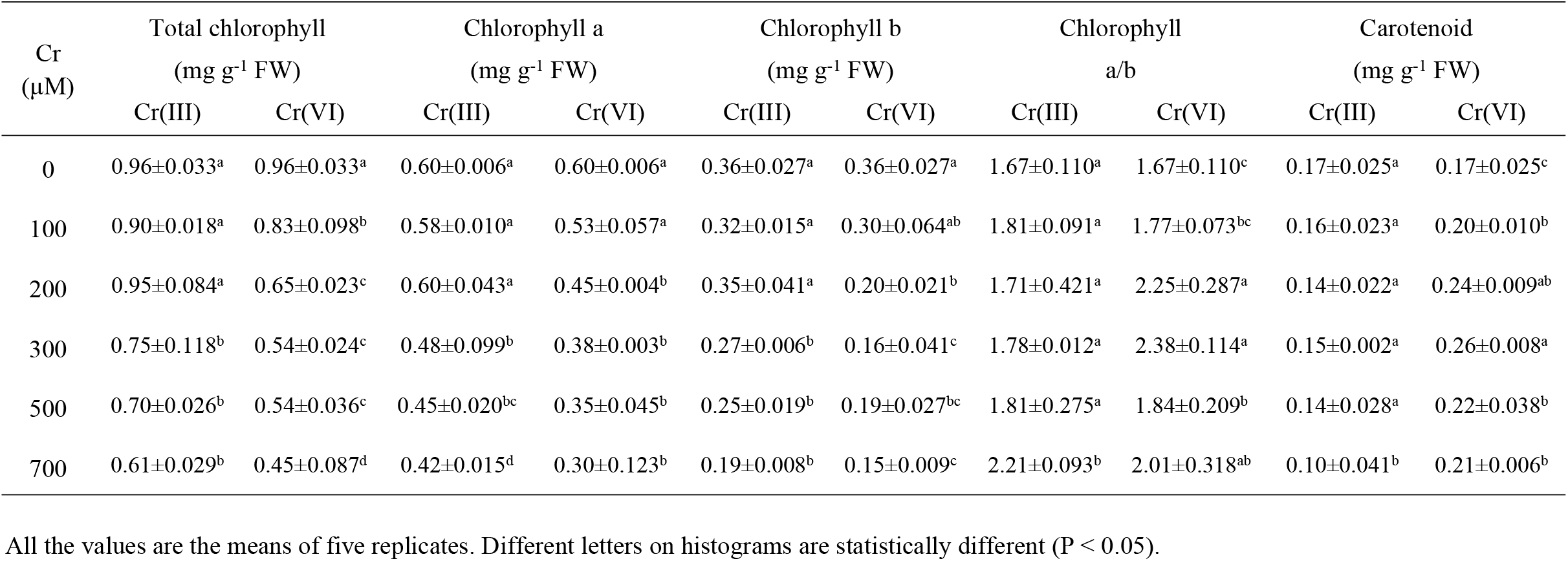
Effects of Cr(III) and Cr(VI) on the content of chlorophyll pigment in the leaves of *A. thaliana*.

But the toxicity caused by Cr(III) and Cr(VI) was different on carotenoid. Carotenoid is a known pigments that protecting plant organs when the plant under stress[28, 29]. In this study, an increase of carotenoid was observed in *A. thalian* treated with Cr(VI). With 300μM of Cr(VI), The carotenoid content increase by 52%. Thus, the continuous increase of carotenoid under Cr(VI) stresses suggest that carotenoid has a role in reducing the toxicity of hexavalent chromium. Cr(VI).

### Antioxidant enzyme analysis

In process of abiotic stresses of plants, Membrane lipids are one of the key cellular targets. The lipid peroxidation is considered to have the function of mobbing free radical[30, 31]. Thus, the contents of lipid peroxidation in roots were determined by the TBA method. All available data suggest that a concentration-dependent increase of MDA in *A. thaliana* roots caused by Cr. The MDA content was higher in *A. thaliana* exposed to Cr(VI) than for Cr(III). (Table 2)

**Table 1.**
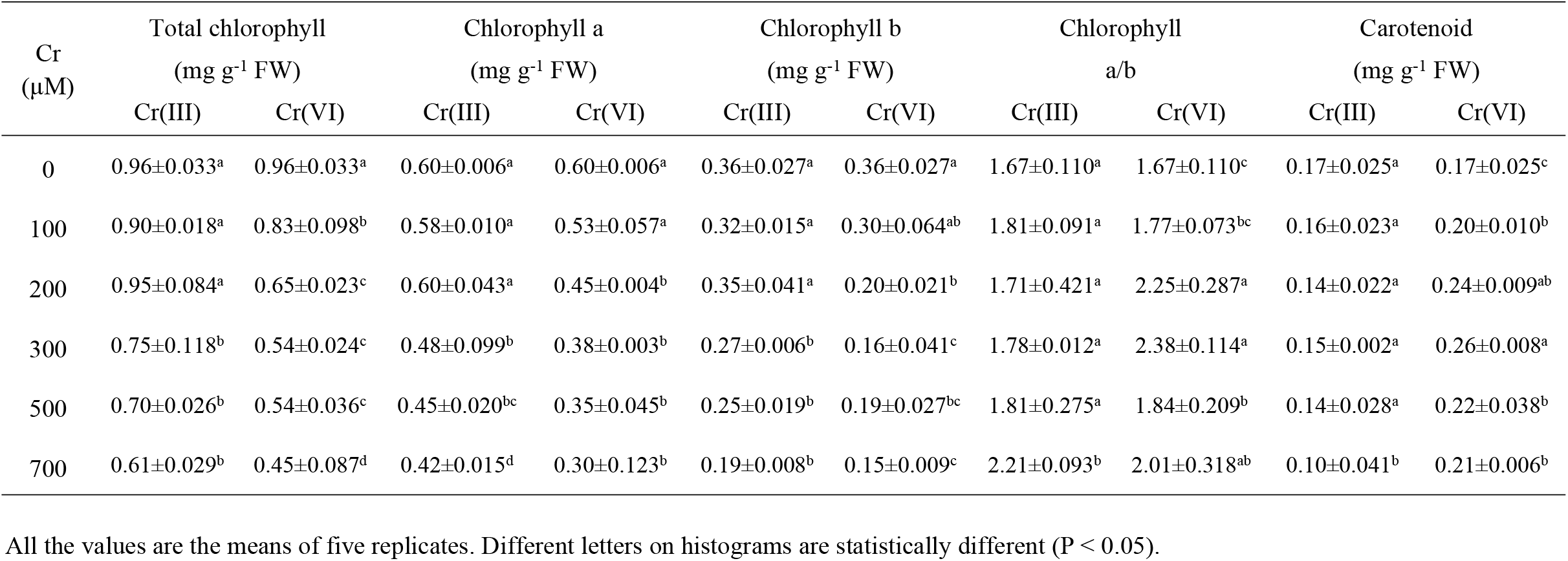
Effects of Cr(III) and Cr(VI) on the content of chlorophyll pigment in the leaves of *A. thaliana*.

In order to mitigate oxidative damage, a series of antioxidant systems were initiate in plants. Practically, antioxidant enzymes such as CAT and SOD are thought to play an important role in the stress process[32, 33]. The POD, CAT, and ascorbate–glutathione cycle (GR and APX), in scavenging H_2_O_2_ are more than important[34]. In this study, a gradual and continual drop of SOD and POD was observed in *A. thalian* treated with Cr(VI) from 100μM and with Cr(III) from 200μM. CAT activity was not significantly affected by Cr stress except treated with 700μM Cr(VI). Chromium stress significantly increases the activity of APX and GR. APX activity enhanced respectively up to 1.7 and 2.1 times of control for the Cr(III) and Cr(VI) at 200μM. With GR, the highest value was 2.1 fold of control at 500μM Cr(III), 2.9 fold at 300μM Cr(VI). It indicates that enzymes engaged in antioxidant defence: APX and GR were activated by Cr, while SOD and POD activity was inhibited.

### The ultra-morphological changes under Cr

Cell mortality of roots under Cr stress was determined by Trypan blue staining. In the results of the trypan blue test, no obvious damage was found in the unstressed blank group. (Fig 3). Staining was detected from 500μM Cr(III) and 200μM Cr(VI), which was more extensive with increasing Cr concentrations.

**Fig. 3.**
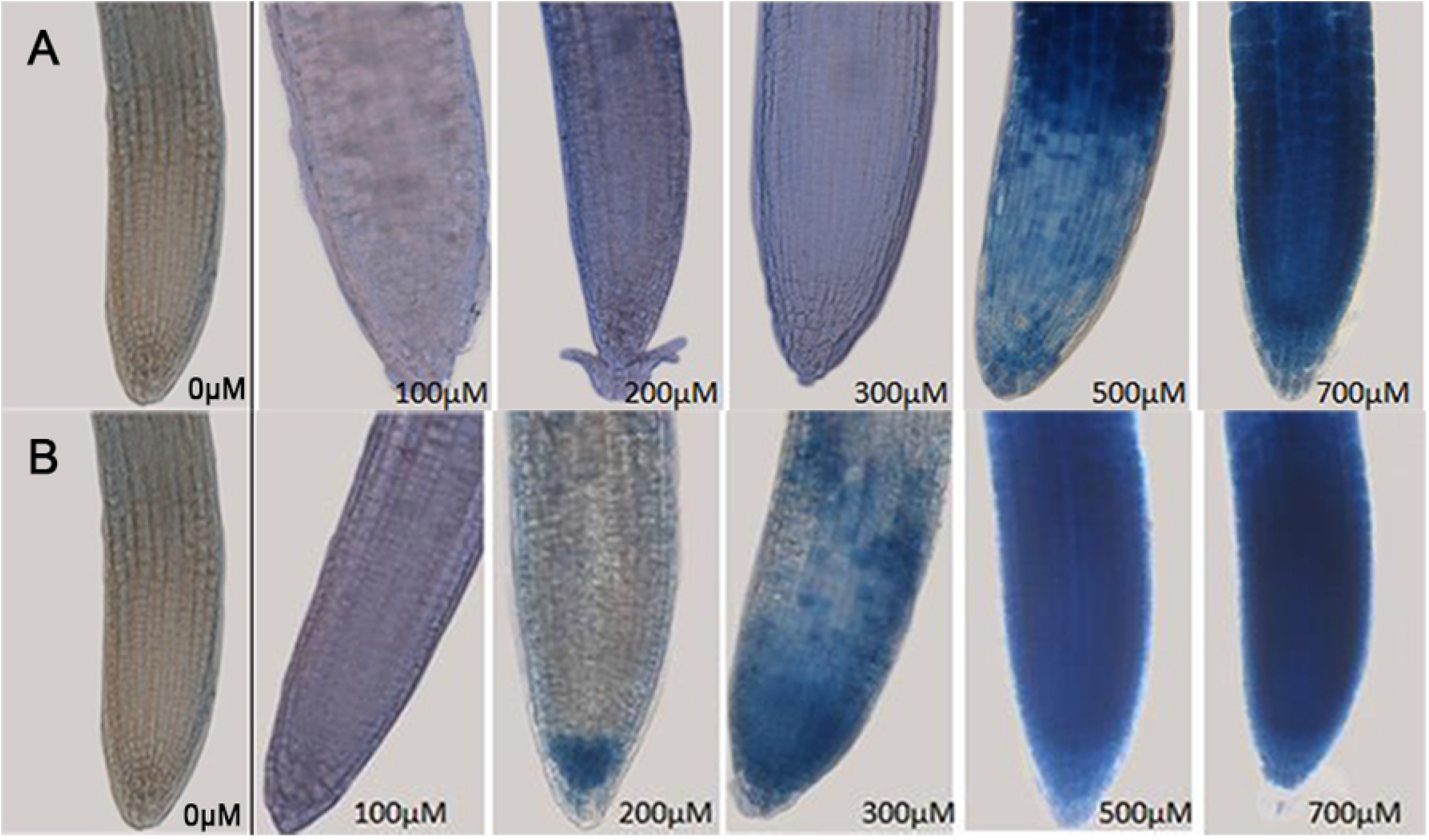
Detection of cell death (by Trypan blue staining) in root tips of A. thaliana. (A) Cr(III) treated roots; (B) Cr(VI) treated roots.

These results of ultra-morphological can be mutually verified by the result of lipid peroxidation experiments. This research shown that the Cr (VI) effect was more heavy than Cr(III)by the results of cell death in roots.

The percentage of intact chloroplasts with orderly arrangement of grana and stroma thylakoid approached 100 percent in the control plant leaves (Fig 4A). Under the Cr(III) treatment, 2-3 starch grains(Sg) appeared and the thylakoid became obscure at 200μM(Fig 4B). At 500μM Cr(III), The continuous increase of Sg is accompanied by further thylakoid disintegrated (Fig 4C). At 200μM Cr(VI), there are 4–6 Sg in chloroplasts and the thylakoids were disorderly arranged (Fig 4D). When the concentration of Cr(VI) reached 500μM, chloroplasts filled with huge Sg, the thylakoid was disappeared(Fig 4E). In conclusion, the chloroplasts of *A. thaliana* can be damaged by Cr(III) or Cr(VI), thus reducing chlorophyll content. This corresponds to the previous result.

**Fig. 4.**
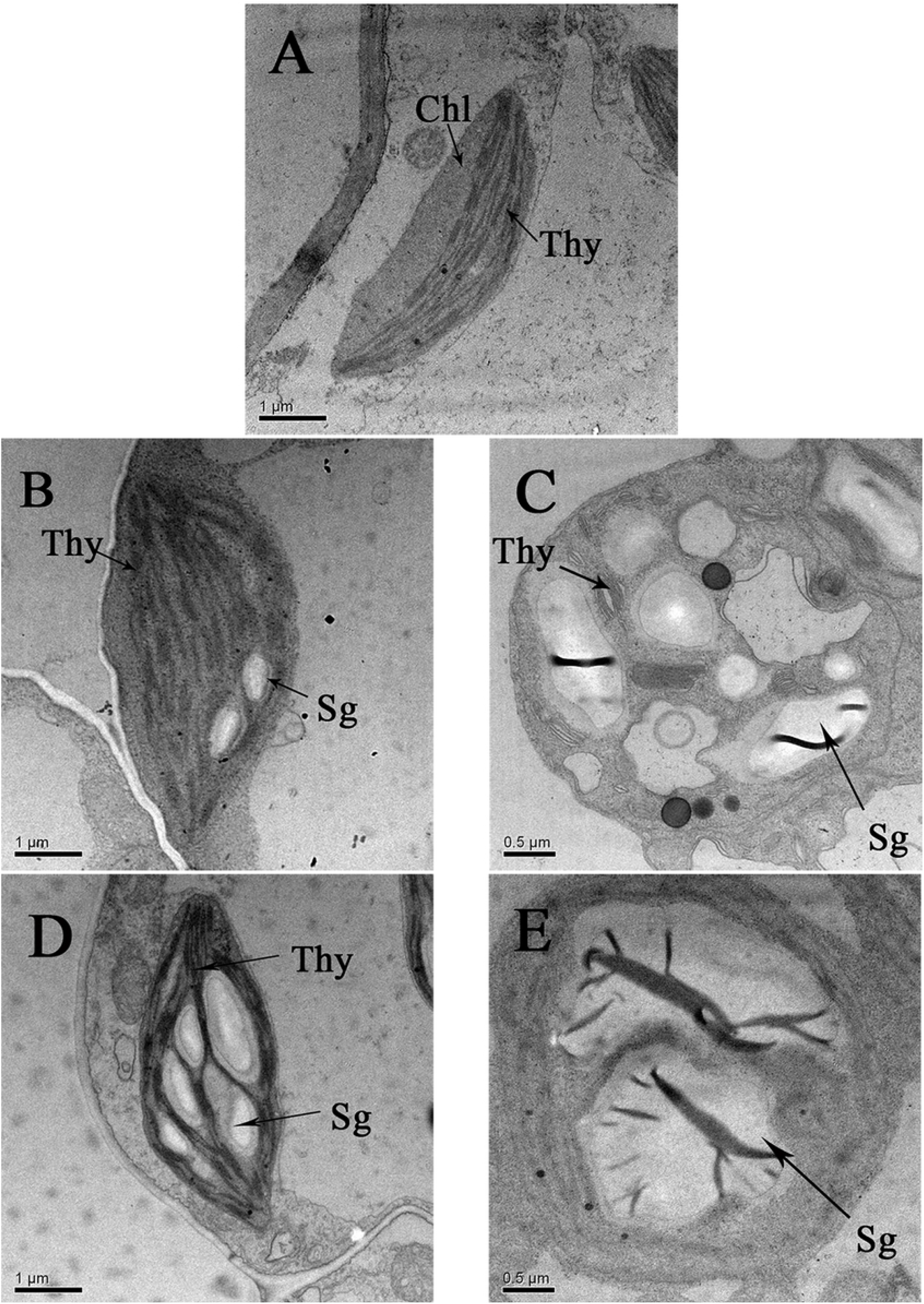
Effects of Cr(III) and Cr(VI) on chloroplasts in leaves’ cells of A. thaliana. (A) Control; (B) Treated with 200μM Cr(III); (C) Treated with 500μM Cr(III); (D) Treated with 200μM Cr(VI) ; (E) Treated with 500μM Cr(VI).

## Conclusions

In this study, we first investigated the physiological, morphological and biochemical reactions of *A. thaliana* under Cr(III) and Cr(VI) stress. The collection capacity for Cr was determined by roots which is the main organ of accumulation, irrespective of Cr(III) and Cr(VI).Our data proved that the intake of Cr(VI) was higher than the intake of Cr(III), in the roots. At 700μM, the Cr(VI) accumulation in the roots of A. thaliana was about 1.5times higher than the Cr(III). Cr toxicity in A. thaliana is closely related to the valence of chromium and the concentration in the nutrient media. It is proved that the toxicity of Cr(VI) is stronger than that of Cr(III) by measuring the growth of A. thaliana. Whether Cr(III) or Cr(VI), when the concentration of Cr reaches or exceeds 200μM, the inhibition of A. thaliana growth was observed. But interestingly, at 200μM, Cr(III) caused a statistically significant increase in dry biomass as compared to the control. It indicated a relatively low concentration of Cr(III) can stimulate A. thaliana growth.

All available data suggest that a concentration-dependent increase of MDA in A. thaliana roots caused by Cr. The MDA content was higher in A. thaliana exposed to Cr(VI) than for Cr(III). These results are consistent with Trypan blue staining assay, and the effect was more pronounced following Cr(VI) exposure. It indicated that A. thaliana suffers from Cr stress, which resulted cell death in roots. For further investigate the mechanism of oxidative stress in Cr toxicity, we focused on some antioxidative enzymes such as SOD, POD, CAT, GR and APX. In this study, a gradual and continual drop of SOD and POD was observed in A. thalian treated with Cr(VI) from 100μM and with Cr(III) from 200μM. CAT activity was not significantly affected by Cr stress except treated with 700μM Cr(VI). Chromium stress significantly increases the activity of APX and GR. APX activity enhanced respectively up to 1.7 and 2.1 times of control for the Cr(III) and Cr(VI) at 200μM. With GR, the highest value was 2.1 fold of control at 500μM Cr(III), 2.9 fold at 300μM Cr(VI). It indicates that enzymes engaged in antioxidant defence: APX and GR were activated by Cr, while SOD and POD activity was inhibited. All data indicated that Cr(VI) was more toxic than Cr(III) in physiological responses.

TEM shows that both Cr(III) and Cr(VI) can cause the chloroplasts damaged and filled the starch grains, chlorophyll a, chlorophyll b and total chlorophyll consequently reduced. But the toxicity caused by Cr(III) and Cr(VI) was different on carotenoid. An increase of carotenoid was measured in A. thalian treated with Cr(VI). With 300μM of Cr(VI), there is an increase of 52%. Thus, consistently increasing amounts of carotenoid under Cr(VI) stresses suggest that Cr(VI) can stimulate the synthesis of carotenoid to detoxified toxicities.

## Acknowledgements

This work was supported by a key scientific and research project grant funded by the Hebei Science and Technology Department (17275505D).

